# Kinase-independent activity of DYRK1A promotes viral entry of highly pathogenic human coronaviruses

**DOI:** 10.1101/2022.09.13.507833

**Authors:** Madison S. Strine, Wesley L. Cai, Jin Wei, Mia Madel Alfajaro, Renata B. Filler, Scott B. Biering, Sylvia Sarnik, Ajinkya Patil, Kasey S. Cervantes, Clayton K. Collings, Peter C. DeWeirdt, Ruth E. Hanna, Kevin Schofield, Christopher Hulme, Silvana Konermann, John G. Doench, Patrick D. Hsu, Cigall Kadoch, Qin Yan, Craig B. Wilen

## Abstract

Identifying host genes essential for severe acute respiratory syndrome coronavirus 2 (SARS-CoV-2) has the potential to reveal novel drug targets and further our understanding of coronavirus disease 2019 (COVID-19). We previously performed a genome-wide CRISPR/Cas9 screen to identify pro-viral host factors for highly pathogenic human coronaviruses. Very few host factors were required by diverse coronaviruses across multiple cell types, but *DYRK1A* was one such exception. Although its role in coronavirus infection was completely unknown, *DYRK1A* encodes <u>D</u>ual Specificity T<u>y</u>rosine Phosphorylation <u>R</u>egulated <u>K</u>inase 1A and regulates cell proliferation, and neuronal development, among other cellular processes. Interestingly, individuals with Down syndrome overexpress *DYRK1A* 1.5-fold and exhibit 5-10x higher hospitalization and mortality rates from COVID-19 infection. Here, we demonstrate that DYRK1A regulates *ACE2* and *DPP4* transcription independent of its catalytic kinase function to support SARS-CoV, SARS-CoV-2, and MERS-CoV entry. We show that DYRK1A promotes DNA accessibility at the *ACE2* promoter and a putative distal enhancer, facilitating transcription and gene expression. Finally, we validate that the pro-viral activity of *DYRK1A* is conserved across species using cells of monkey and human origin and an *in vivo* mouse model. In summary, we report that DYRK1A is a novel regulator of *ACE2* and *DPP4* expression that may dictate susceptibility to multiple highly pathogenic human coronaviruses. Whether DYRK1A overexpression contributes to heightened COVID-19 severity in individuals with Down syndrome through *ACE2* regulation warrants further future investigation.

## INTRODUCTION

Severe acute respiratory syndrome coronavirus 2 (SARS-CoV-2), the causative agent of coronavirus disease 2019 (COVID-19), is a beta coronavirus that has launched an ongoing pandemic and continues to threaten public health globally^1,2^. Two additional beta coronavirus family members (SARS-CoV and Middle East respiratory syndrome coronavirus (MERS-CoV)) have caused more limited epidemics but with a higher case fatality rate^3–5^. There are also four endemic alpha and beta human coronaviruses cause the common cold (HCoV-NL63, HcoV-OC43, HcoV-229E, and HcoV-HKU1). Despite zoonotic outbreaks of three beta coronaviruses in less than 20 years, our understanding of host factors that support these highly pathogenic human coronaviruses remain incompletely understood^1,6^.

The coronavirus life cycle commences with viral entry which requires receptor binding and subsequent proteolytic processing of the viral spike (S) glycoprotein^7^. For initial infection and viral spread, a variety of epithelial cells can be targeted by SARS-CoV-2, but ciliated cells are the predominant target^8,9^. SARS-CoV and SARS-CoV-2 coronaviruses use angiotensin-converting enzyme 2 (ACE2) receptor, whereas MERS-CoV engages dipeptidyl peptidase-4 (DPP4) as a receptor^10–14^. After receptor binding, the spike glycoprotein undergoes proteolytic cleavage at the cell surface by the transmembrane serine protease 2 (TMPRSS2) or in the endosome by Cathepsin L (CTSL), enabling cell surface-mediated or endosomal entry, respectively^12,15–18^. Once internalized and proteolytically primed, the viral and host membranes fuse releasing the viral RNA genome and enabling viral protein translation and establishment of viral replication complexes^19–21^. Viral structural proteins (nucleocapsid, spike, membrane, and envelope) package the nascent viral genome genomes into mature virions, which egress from the cell enabling viral spread^19–21^.

We performed a genome-wide CRISPR/Cas9-based inactivation screen in African green monkey kidney Vero-E6 cells and in human lung epithelial Calu-3 cells^22,23^. Both Vero-E6 and Calu-3 cells are permissive to SARS-CoV, SARS-CoV-2, and MERS-CoV and endogenously express the cognate receptors ACE2 and DPP4^22–26^. Our screens revealed several pro-viral genes shared by SARS-CoV, SARS-CoV-2, and MERS-CoV in both Vero-E6 and Calu-3 cells^22,23^. Other genome-wide CRISPR/Cas9 screens for SARS-CoV-2 and other related coronaviruses have since identified additional host dependency factors including proteins involved in viral entry, endocytic trafficking and sorting, cholesterol homeostasis, and autophagy^27–32^. While reproducibility between screens was excellent when performed on the same cell type, there was otherwise limited overlap across hits in these loss of function screens^27–32^. One notable exception was DYRK1A, which was a top enriched hit in both Vero-E6 and Calu-3 cells^22,23,33,34^. Interestingly, DYRK1A was not a hit in cell lines exogenously overexpressing ACE2, such as A549, Huh7.5, or HeLa cells^27–32^. Here, we sought to elucidate how DYRK1A regulates CoV infection and to determine the cell-type specificity of its function. Because DYRK1A was identified in cells with high endogenous ACE2 expression and was also a top hit for a replication competent SARS-CoV-2 pseudovirus, we posited that DYRK1A may function as a novel transcriptional regulator of coronavirus entry^22^. Preventing or reducing viral entry can circumvent the downstream processes of the coronavirus lifecycle, including pathogenesis and spread. While SARS-CoV-2 mRNA vaccines have proved highly efficacious at achieving this, the mechanisms that govern receptor expression represent a major gap in our knowledge of coronavirus biology^35^.

*DYRK1A* encodes <u>D</u>ual Specificity T<u>y</u>rosine Phosphorylation <u>R</u>egulated <u>K</u>inase 1A, a member of the CMGC kinase group that includes cyclin-dependent kinases, mitogen-activated protein kinases, glycogen synthase kinases, and CDC-like kinases^36,37^. DYRK1A encodes a bipartite nuclear localization sequence and, as a result, accumulates predominantly in the nucleus, although cytosolic DYRK1A is reported in some contexts^38,39^. Located on chromosome 21 in the Down syndrome critical region (21q22.22), *DYRK1A* is highly dosage-sensitive gene^40–42^. In Down syndrome (also known as trisomy 21), there is an extra copy of *DYRK1A* resulting in overexpression that is strongly associated with disease pathogenesis and neurological developmental delay^43–47^. In contrast, downregulated expression of *DYRK1A* can cause haploinsufficiency syndromes associated with microcephaly and autism spectrum disorder^48,49^. Interestingly, individuals with trisomy 21 are highly susceptible to SARS-CoV-2 with significantly elevated (five- to ten-fold) risk of infection, hospitalization, and death^49–54^. With such an increase in morbidity and mortality, Down syndrome may among the top genetic disorders associated with the highest risk for COVID-19. The mechanisms underlying this increased COVID-19 morbidity and mortality are unknown.

DYRK1A shares notable identity with its *Drosophila* ortholog Minibrain and is highly conserved across lower to higher eukaryotes^36,55,56^. Regulation of DYRK1A is tightly controlled but poorly understood and modulates a myriad of functions including transcription, protein localization, protein-protein interactions, and proteolytic degradation^36,37,57–60^. Autophosphorylation of the DYRK1A tyrosine residue Y^321^ causes irreversible catalytic Dyrk1a activation^37,59,61–63^. Once activated, DYRK1A acts as a mature serine/threonine kinase, with multiple phospho-substrates involved in cell cycle regulation, development, cell-cell signaling, and transcription, among others^36,37,56^. DYRK1A also has kinase-independent functions, including mRNA stabilization and transcriptional activation^64–68^. Mechanistic insights into these catalytically independent functions are lacking, but data suggests DYRK1A may operate as a scaffold in these cases^39,65,69^.

Despite its roles in many biological processes, the role of DYRK1A in viral pathogenesis remains incompletely explored. DYRK1A can promote Human Papillomavirus Type 16 infection by phosphorylating the viral oncoprotein E7^70^. Similarly, DYRK1A enables transformation by interacting with the oncoprotein E1A from human adenovirus type 5 (HAdV-5)^71^. Inhibition of DYRK1A kinase activity or genetic knockdown of DYRK1A also reduces human cytomegalovirus (HCMV) replication^72,73^. In the cases of both HCMV and HAdV-5, how DYRK1A performs this function is unclear. DYRK1A also possesses anti-viral roles, such as by reducing HIV replication *in vitro* by downregulating cyclin L2 and inhibiting long terminal repeat-driven transcription^74,75^.

Here, we show DYRK1A is critical for highly pathogenic human coronavirus infection *in vitro* and *in vivo*. We demonstrate DYRK1A is a novel regulator of coronavirus entry for both SARS-CoVs and MERS-CoV by promoting ACE2 and DPP4 receptor expression at the mRNA level. We reveal that DYRK1A performs its pro-viral role in the nucleus independently of kinase function, suggesting a previously undescribed mechanism of DYRK1A activity in viral infection.

## RESULTS

### DYRK1A promotes viral entry for SARS-CoVs and MERS-CoV

Our group and others recently identified DYRK1A as a critical host factor for SARS-CoV-2, MERS-CoV, and chimeric HKU5 (bat coronavirus) expressing the SARS-CoV-1 spike (HKU5-SARS-CoV-1-S) in a genome-wide CRISPR/Cas9 inactivation screen in Vero-E6 cells^22,33^. In two additional independent genome-wide CRISPR screens in Calu-3 human immortalized lung cancer cells, DYRK1A was also identified as a top pro-viral gene for SARS-CoV-2^23,33^. By comparing the top 10,000 enriched genes ranked by z-score for SARS-CoV-2 across Vero-E6 and Calu-3 cells, we identified *DYRK1A* as second most strongly enriched gene after only *ACE2*^22,23^ (**Fig. 1A**). In another screen, *DYRK1A* was the third most strongly enriched hit after *ACE2*^33^. We generated single-cell knockout (KO) cells of DYRK1A in Vero-E6 cells using CRISPR/Cas9 to validate screening results and clarify the pro-viral mechanism underlying DYRK1A activity in coronavirus infection (**Fig. 1B**). We challenged clonal KO cells with an infectious clone of SARS-CoV-2 encoding the fluorescent reporter mNeonGreen (icSARS-CoV-2mNG) and quantified viral infection by microscopy^76^. Cells deficient in DYRK1A were resistant to SARS-CoV-2 infection (**Fig. 1C**). Consistent with this finding and our screening results, loss of DYRK1A conferred resistance to virus-induced cell death by SARS-CoV-2, HKU5-SARS-CoV, and MERS-CoV (**Fig. 1D**). We next tested whether DYRK1A acts at viral entry using a pseudotype assay, where coronavirus spike proteins are expressed on a replication deficient vesicular stomatitis virus (VSV) encoding a luciferase reporter. This pseudovirus enables a single round of spike-dependent viral entry. Like loss of ACE2 or CTSL, cells lacking DYRK1A exhibited a significant defect in viral entry for pseudoviruses expressing CoV spike proteins (**Fig. 1E; Extended Data Fig. 1**). These data highlight DYRK1A as a pro-viral host factor for these highly pathogenic human coronaviruses, including SARS-CoV-2, that regulates spike-mediated viral entry in Vero-E6 cells.

**Figure 1.**
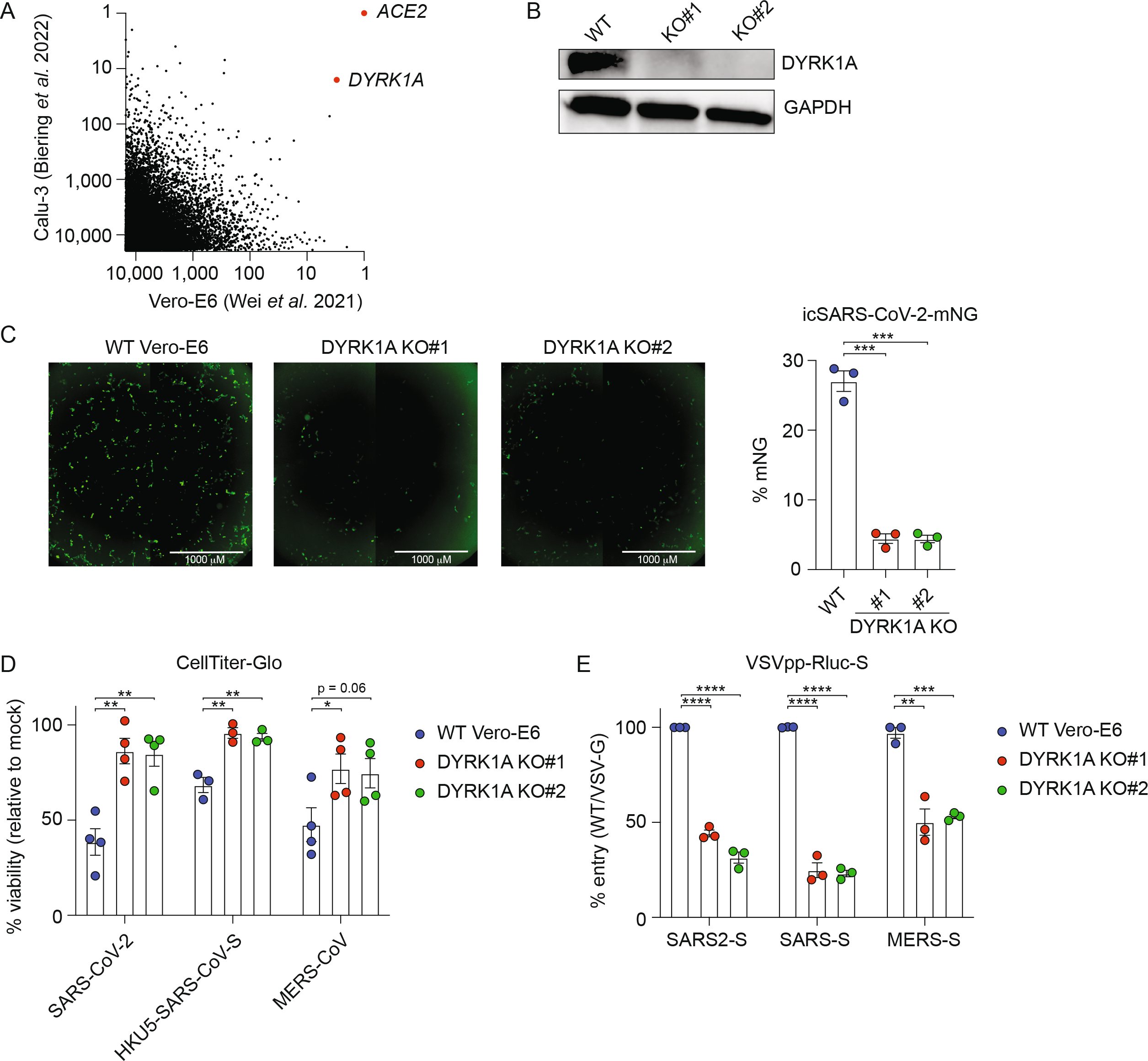
DYRK1A promotes viral entry for SARS-CoV, SARS-CoV-2, and MERS-CoV. **(A)** XY plot comparing the top 10,000 enriched genes that promote SARS-CoV-2 infection in genome-wide CRISPR screens performed in Wei *et al*. 2021 (Vero-E6 cells, African green monkey kidney cells) and Biering *et al*. 2022 (Calu-3 cells, human lung epithelial cells). DYRK1A scored as the most strongly enriched gene after ACE2, supporting a conserved pro-viral role for DYRK1A in monkey and human cells. **(B)** Immunoblot for two single-cell monoclonal knockouts (KO) of DYRK1A (KO#1 and KO#2) generated from parental Vero-E6 cells. Cells lacking DYRK1A were confirmed by Sanger sequencing. **(C)** Wild-ype (WT) Vero-E6 cells and DYRK1A KO cells were infected with icSARS-CoV-2-mNeonGreen (mNG) at an MOI ~ 1.0 and imaged at 48 hours post-infection (hpi) (left). mNeonGreen+ expressing cell frequency (%mNG) was quantified from stitched images (right). Scale bar: 1000 μm. **(D)** DYRK1A KO and WT Vero-E6 cells were infected with SARS-CoV-2, HKU5-SARS-CoV-S, or MERS-CoV at an MOI ~ 1.0 and cell viability was assessed at 72 hpi with CellTiter Glo. % Viability was calculated relative to uninfected controls. **(E)** DYRK1A KO and WT Vero-E6 cells were infected with VSV pseudovirus (VSVpp) encoding CoV spike proteins and a Renilla luciferase (Rluc) reporter at 24 hpi. % Entry for VSVpp-Rluc-SARS2-S, VSVpp-Rluc-SARS-S, and VSVpp-Rluc-MERS-S was normalized to VSVpp-Rluc-VSV-G control and WT Vero-E6 cells. Data were analyzed by unpaired Student’s t-test; * p< 0.05, ** p< 0.01, *** p< 0.001, **** p< 0.001. Shown are means ± SEM. Data in **(C)**, **(D)**, and **(E)** are representative of three independent biological experiments performed with at least 3 technical replicates.

### DYRK1A regulates expression of the receptors ACE2 and DPP4 at the transcript level

Because coronavirus entry is receptor dependent, we next compared the expression of the receptors ACE2 and DPP4 in wild-type cells and DYRK1A KO cells. In DYRK1A KO cells, ACE2 expression is notably reduced at the protein level by Western blot (**Fig. 2A)** but endogenous DPP4 expression was too low to detect even in wild-type cells (**Fig. 2D**, left). To determine whether DYRK1A regulates ACE2 or DPP4 expression at the mRNA level, we performed RT-qPCR to quantify mRNA abundance (**Fig. 2B**). Loss of DYRK1A causes a significant reduction in mRNA transcript levels of both *ACE2* and *DPP4*, suggesting that DYRK1A may transcriptionally regulate the *ACE2* and *DPP4* loci or that DYRK1A may modulate the mRNA stability of these transcripts (**Fig. 2B**). We then overexpressed *ACE2* or *DPP4* in *DYRK1A* KO cells by stable overexpression to clarify whether post-entry or protease-dependent entry may also be regulated by DYRK1A (**Fig. 2C-D**). Overexpression of ACE2 and DPP4 both rescued the entry defect conferred by loss of DYRK1A for SARS-CoVs and MERS-CoV pseudoviruses, respectively (**Fig. 2C-D**). Taken together, these data indicate that DYRK1A promotes viral entry by supporting the expression of *ACE2* and *DPP4* at the mRNA level and that this regulation is independent of post-entry.

**Figure 2.**
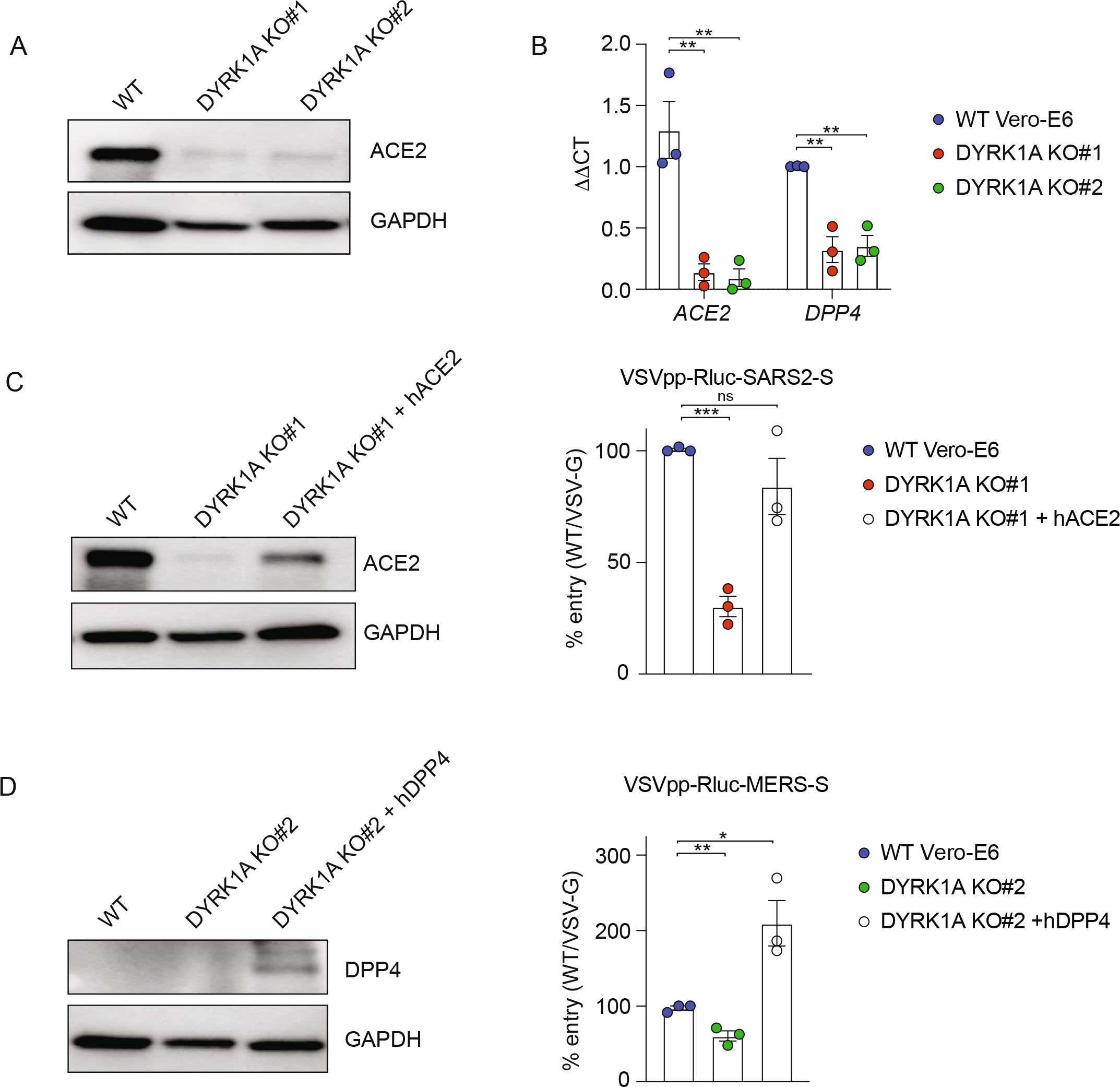
DYRK1A regulates expression of ACE2 and DPP4. **(A)** Immunoblot for ACE2 in WT Vero-E6 and DYRK1A KO cells. **(B)** mRNA abundance of ACE2 and DPP4 transcripts in WT Vero-E6 and DYRK1A KO clones assessed by qPCR. ddCT values are calculated relative to actin and normalized to WT Vero-E6 values. **(C)** WT Vero-E6, DYRK1A KO#1, and DYRK1A KO#1 overexpressing recombinant human ACE2 (hACE2) were infected with VSVpp-Rluc-SARS2-S and % entry was assessed at 24 hpi. Immunoblot confirms rescue of ACE2 expression in DYRK1A KO#1. **(D)** WT Vero-E6, DYRK1A KO#2, and DYRK1A KO#2 overexpressing recombinant human DPP4 (hDPP4) were infected with VSVpp-Rluc-MERS-S and % entry was assessed at 24 hpi. Immunoblot confirms DPP4 overexpression in DYRK1A KO#2. % Entry in **(C)** and **(D)** was normalized to VSVpp-Rluc-VSV-G and WT Vero-E6 cells. Data were analyzed by unpaired Student’s t-test; ns p> 0.05, * p< 0.05, ** p< 0.01, *** p< 0.001. Shown are means ± SEM. Each datapoint is the mean of 3-5 technical replicates. All experiments were performed in biological replicate.

### The pro-viral role of DYRK1A is kinase-independent in nature

To confirm these pro-viral phenotypes and to elucidate the mechanism of DYRK1A pro-viral activity, we reintroduce DYRK1A into our single-cell KO clones (**Fig. 1B**) by lentiviral transcription. We transduced the following constructs: wild-type DYRK1A, kinase-null DYRK1A (encoding K188R or Y321F inactivating point mutations), and nuclear localization mutant DYRK1A (encoding a nuclear export signal and disruption of the bipartite nuclear localization motif) (**Fig. 3A**). The active site of DYRK1A requires ATP for catalytic activity, and DYRK1A phosphorylation activity is inhibited by preventing ATP binding with Lys^188^ by mutating the residue to arginine (K188R)^61,62,77^. Disruption of Y^321^ autophosphorylation by mutating tyrosine to phenylalanine (Y321F) renders DYRK1A catalytically inactive by preventing kinase maturation^61,78–80^. Because sustained DYRK1A overexpression can cause cell cycle exit^81–83^, we generated these constructs under the control of a doxycycline inducible Tet-on promoter. We first confirmed that these addbacks could rescue DYRK1A and ACE2 expression (**Fig. 3B**). Consistent with previous literature, expression of DYRK1A-K188R is weaker relative to wild-type and DYRK1A-Y321F constructs, likely due to protein destabilization (**Fig. 3B**)^77^. Complementation with wild-type DYRK1A restored ACE2 expression (**Fig. 3B**). Unexpectedly, loss of kinase activity (K188R and Y321F) enabled at least partial rescue of ACE2 expression by Western blot, whereas loss of nuclear localization did not elicit detectable ACE2 protein expression, despite partial retention of DYRK1A in the nucleus (**Fig. 3B-C**). To confirm DYRK1A complementation could rescue infection with authentic virus, we challenged these cell lines with the icSARS-CoV-2-mNG reporter virus^76^ (**Fig. 3D**). Consistent with protein expression data, wild-type and kinase-dead constructs significantly rescued infection in DYRK1A KO cells. Interestingly, the nuclear localization mutant DYRK1A weakly rescued infection, possibly due to residual DYRK1A in the nucleus (**Fig. 3B, D**). Next, we tested whether full or partial restoration of DYRK1A and ACE2 expression could rescue viral entry. In all cases where DYRK1A was reintroduced, DYRK1A KO clones exhibited full or partial rescue of SARS-CoV-2 and MERS-CoV viral entry, including catalytic and localization mutants (**Fig. 3E**). Because genetic approaches indicated that DYRK1A kinase activity is dispensable for SARS-CoV-2 infection, we next decided to validated these findings using a pharmacologic approach. We inhibited DYRK1A with harmine, INDY, DYR219, or DYR533 – four potent and specific type I ATP-competitive kinase inhibitors (**Fig. 3F**)^84–88^. Pharmacologic inhibition of DYRK1A had no effect on SARS-CoV-2 virus-induced cell death, in contrast to a protease inhibitor (calpain inhibitor III), confirming our genetic findings (**Fig. 3F**). Collectively, these data support that nuclear DYRK1A regulates ACE2 expression in a kinase-independent manner.

**Figure 3.**
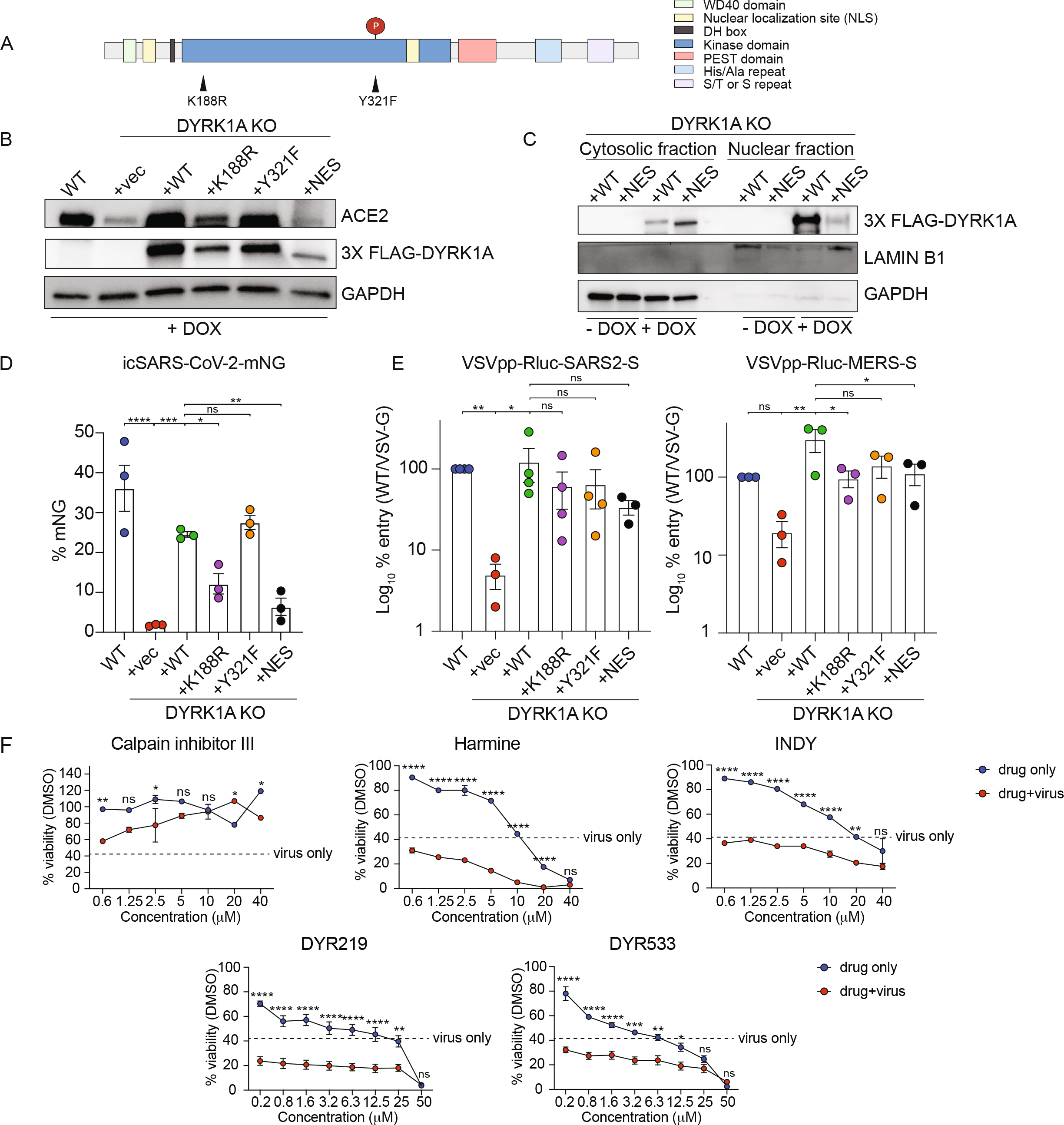
The pro-viral role of DYRK1A is kinase-independent. **(A)** DYRK1A protein domains and engineered point mutations. Nuclear localization mutants were generated by deletion of the bipartite nuclear localization motif and addition of a C-terminal nuclear export signal. **(B)** Immunoblot for cells expressing an empty vector (KO+vec), WT DYRK1A (KO+WT), kinase dead DYRK1A (KO+K188R and KO+Y321F), or a nuclear localization mutant DYRK1A (KO+NES) were reintroduced to DYRK1A KO#1. Constructs were tagged with 3X FLAG and induced by doxycycline (DOX) for 72 hours. Reintroduction of DYRK1A rescues ACE2 expression was rescued relative to WT Vero-E6 cells. **(C)** Immunoblot after cytosolic-nuclear fractionation of DYRK1A KO+WT or DYRK1A KO+NES, demonstrating that DYRK1A is predominantly localized to the nucleus until disruption of the bipartite nuclear localization motif and addition of a nuclear export signal. DYRK1A expression was induced by DOX for 72 hours prior to fractionation. **(D)** WT Vero-E6 cells and cells overexpressing DYRK1A after DOX induction for 72 hours were infected with icSARS-CoV-2-mNeonGreen (mNG) at MOI ~ 1. Cells were imaged and mNeonGreen+ expressing cell frequency (%mNG) was quantified from stitched images. **(E)** WT Vero-E6 cells and DYRK1A KO cells with reintroduced DYRK1A were infected with VSVpp-Rluc-SARS2-S or VSVpp-Rluc-MERS-S and % entry was assessed at 24 hpi. % Entry was normalized to VSVpp-Rluc-VSV-G and WT Vero-E6 cells. **(F)** WT Vero-E6 cells were treated with the positive control protease inhibitor calpain inhibitor III or a potent DYRK1A inhibitor (harmine, INDY, DYR219, and DYR533) for 48 hours. Cells were then infected with SARS-CoV-2 (MOI ~ 1) and cell viability was assessed 72 hpi via CellTiter Glo. % Viability was calculated relative to uninfected or untreated controls. Data were analyzed by Kruskal-Wallis (C, left), ordinary one-way ANOVA **(D, E** left), or two-way ANOVA multiple comparisons tests **(E** right); ns p> 0.05, * p< 0.05, ** p< 0.01, *** p< 0.001, **** p< 0.0001. Data shown are means ± SEM of **(D, E)** 5-10 technical replicates across three independent experiments and **(F)** 2-3 technical replicates performed in biological duplicate.

### DYRK1A drives ACE2 and DPP4 expression by altering chromatin accessibility

Because DYRK1A can positively regulate transcription and gene expression^58,64,65,67,68^, we next profiled global gene expression (RNA-seq) and chromatin accessibility (ATAC-seq) to assess the mechanism of DYRK1A-mediated regulation of *ACE2* and *DPP4* (**Fig. 4**; **Fig. 2S**). Loss of DYRK1A resulted in up- or down-regulation of approximately ~2,000 genes. Analysis of differentially expressed genes (DEGs) confirmed that loss of DYRK1A confers downregulation of *ACE2* and *DPP4*, as well as *CTSL*, which encodes the protease for spike cleavage and viral entry in Vero-E6 cells but not Calu-3 cells^24^ (**Fig. 4B-C; Extended Data Fig. 2A**). Importantly, complementation with both DYRK1A-WT and DYRK1A-Y321F rescue expression of these genes (**Fig. 4B, D**). There was no correlation between DYRK1A-dependent gene regulation and CRISPR genes (ranked by z-score) with the exception of ACE2, DPP4, and CTSL^22^ (**Fig. 4C**). Gene ontology analysis also revealed little overlap between biological pathways regulated by DYRK1A and those involved in SARS-CoV-2 infection (**Extended Data Fig. 2B**)^22^. Because RNA-seq revealed that DYRK1A promotes increased mRNA levels for these genes, we next asked whether DYRK1A altered chromatin accessibility at these sites, thereby altering transcription. Loss of DYRK1A confers generally more open chromatin states across the genome (**Extended Data Fig. 2C**). In contrast, absence of DYRK1A resulted in reduced accessibility near the *ACE2* transcriptional start site (TSS), a putative proximal enhancer, and a putative distal enhancer situated within *BMX* (**Fig. 4F; Extended Data Fig. 2C**). (Wei and Patil *et al*. 2022). Chromatin accessibility was restored at these sites upon reintroduction of DYRK1A (**Fig. 4F; Extended Data Fig. 2C**). A second putative distal enhancer located within *ASB11*, does not seem to be altered by DYRK1A (p > 0.1), indicating that DYRK1A-mediated regulation of *ACE2* may be context- or site-specific (**Fig. 4F**). Interestingly, three sites were identified where loss of DYRK1A led to significantly increased (p < 0.0005) chromatin accessibility within 5 kb of the *ACE2* TSS, suggesting that DYRK1A may also close off chromatin via repressive activity at the *ACE2* locus (**Extended Data Fig. 2C**). In comparison to SMARCA4, another *ACE2* regulator we have identified, DYRK1A-mediated DNA accessibility significantly overlaps with SMARCA4 at approximately 1/3 of sites (p<0.00001 by Fisher’s exact t-test), suggesting that some pathways may be co-regulated by the two (**Extended Data Fig. 3D**). However, DYRK1A-mediated DNA accessibility (correlation coefficient ~ 0.33) and gene expression output (correlation coefficient ~ 0.08) overall poorly correlate with those regulated by SMARCA4 (**Extended Data Fig. 3C, E**) (Wei and Patil *et al*. 2022). Moreover, SMARCA4 promotes open chromatin states at both putative distal enhancers (*BMX* and ASB11) and the putative *ACE2* proximal promoter (Wei and Patil *et al*. 2022). Together, these data suggest that DYRK1A regulates the majority of DNA accessibility and gene expression independently of SMARCA4. Therefore, our data supports that DYRK1A is a critical regulator of *ACE2* chromatin accessibility and transcription by a novel mechanism. Although enhancer regions and other regulatory elements are not well defined for *DPP4* and *CTSL*, we observed increased chromatin accessibility at sites proximal and distal to the respective TSS for these genes, which was not the case for SMARCA4 (**Fig. 4G-H**). Unlike in the case of *ACE2*, putative insulator regions (i.e., sites where loss of DYRK1A led to more open chromatin) were not identified for *DPP4* or *CTSL*. These data highlight that DYRK1A – albeit not a canonical epigenetic modifying enzyme or transcription factor – can dramatically alter chromatin accessibility to drive transcription of a pro-viral gene expression axis that promotes viral entry.

**Figure 4.**
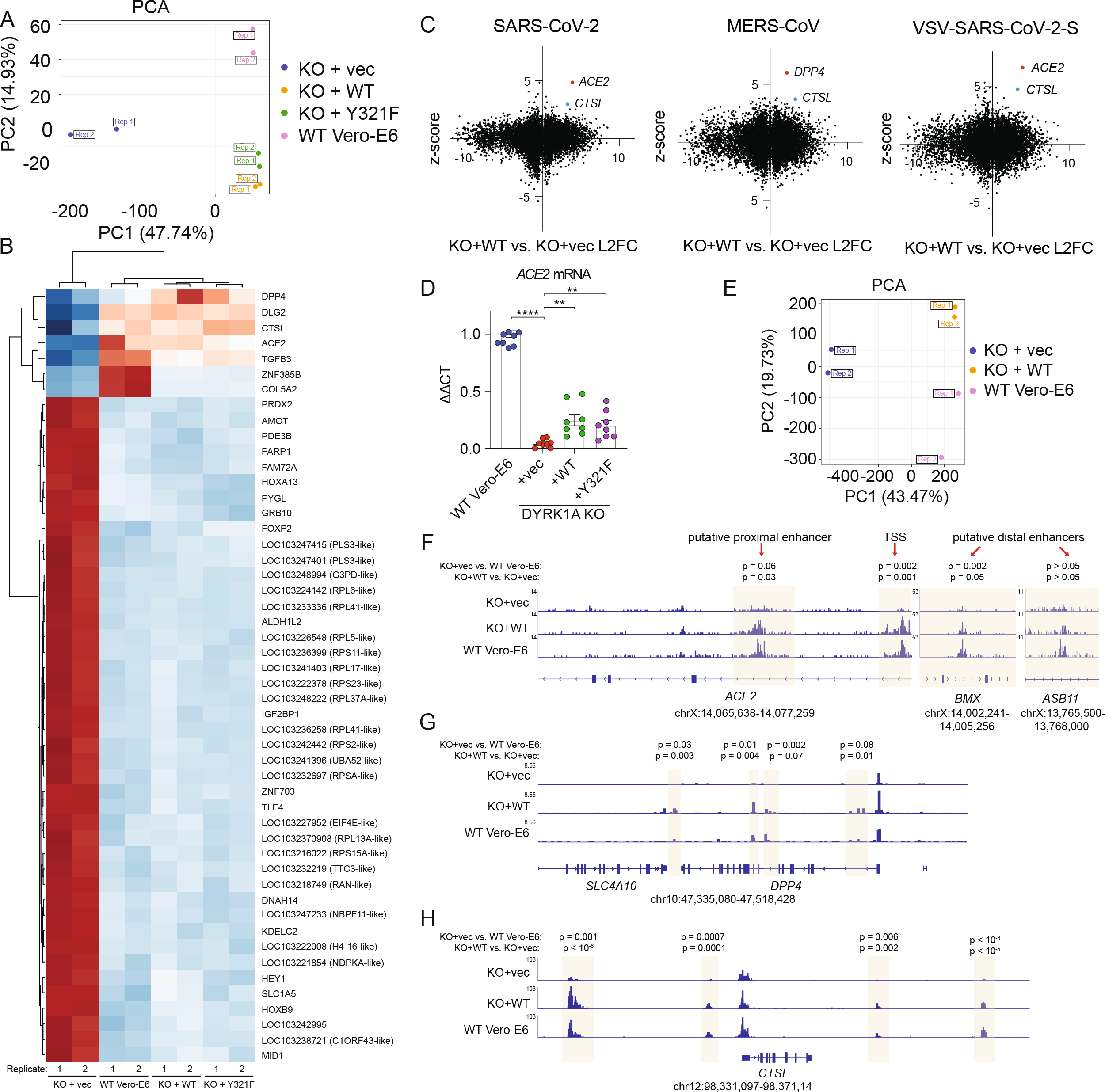
DYRK1A drives ACE2 and DPP4 expression by altering chromatin accessibility. **(A)** Principal component analysis of RNA-Seq experiments performed in WT Vero-E6, KO+vec, KO+WT, and KO+Y321F cells. Rep refers to independent biological replicates. **(B)** Heatmap depicting differentially expressed genes by RNA-Seq in WT Vero-E6, KO+vec, KO+WT, and KO+Y321F cells. **(C)** XY plot of RNA-Seq L2FC versus CRISPR z-scores in Vero-E6 cells (Wei *et al*. 2022) for SARS-CoV-2 (left), MERS-CoV (middle), or VSV-SARS-CoV-2-S (right). Denoted are receptor (*ACE2, DPP4*) and protease (*CTSL*) genes as significantly upregulated pro-viral genes of interest. **(D)** qPCR for *ACE2* validates RNA-Seq results and supports partial rescue of *ACE2* mRNA transcripts in cells where DYRK1A-WT or DYRK1A-Y321F are reintroduced. **(E)** Principal component analysis of ATAC-Seq experiments performed in WT Vero-E6, KO+vec, KO+WT, and KO+Y321F cells. Rep refers to independent biological replicates. **(F)** ATAC-Seq gene tracks for *ACE2*, highlighting increased accessibility at putative enhancers and near the transcriptional start site (TSS) in the presence of DYRK1A. **(G)** ATAC-Seq gene tracks for *DPP4*, showing increased chromatin accessibility in the presence of DYRK1A. **(H)** ATAC-Seq genome tracks for *CTSL*, showing increased chromatin accessibility in the presence of DYRK1A. All experiments were performed in biological duplicate (RNA-seq/ATAC-seq) or triplicate (qPCR).

### DYRK1A is a conserved pro-viral factor for SARS-CoV-2 in human lung epithelial cells and in mice

We then sought to determine whether the pro-viral role of DYRK1A observed in Vero-E6 cells was conserved in human cells and murine models of SARS-CoV-2 infection. We previously performed a subpool screen in Calu-3 cells that validated DYRK1A as a positive regulator of SARS-CoV-2 infection^22^. To validate the results of that screen, we generated pooled knockouts in Calu-3 cells using guides targeting *DYRK1A* or a non-targeting guide control. In Calu-3 cells, loss of *DYRK1A* causes a significant reduction in SARS-CoV-2 infection assessed by Tissue Culture Infectious Dose (TCID_50_) assay (**Fig. 5A**). Next, we tested whether DYRK1A could promote SARS-CoV-2 infection *in vivo* using the mouse adapted SARS-CoV-2 strain MA10^89^. Since loss of DYRK1A is embryonically lethal^90^, we crossed an existing DYRK1A conditional deletion mouse (*DYRK1A*^F/F^) to a tamoxifen-inducible Cre recombinase under control of the globally expressed ubiquitin c gene (*Ubc* CreERT2) (**Fig. 5C**). In parallel, we also generated a conditional knockout for ACE2 in a human ACE2 overexpressing mouse (*hACE2* KI/het x *Ubc* CreERT2) to use as a positive control (**Fig. 5B**). We treated mice with tamoxifen for five days to conditionally ablate DYRK1A and then infected mice with 10^5^ plaque forming units (PFU) of MA10 or SARS-CoV-2 intranasally. One day post-infection, mice were sacrificed and lungs were harvested and homogenized to assess viral load by plaque assay. Loss of both *hACE2* and *DYRK1A* significantly reduced viral titers in lungs, supporting a pro-viral role for DYRK1A *in vivo* (**Fig. 5B-C**). Overall, these findings suggest that DYRK1A is a critical host factor for SARS-CoV-2 infection that is conserved across monkey (Vero-E6) cells, human lung cells (Calu-3), and mice.

**Figure 5.**
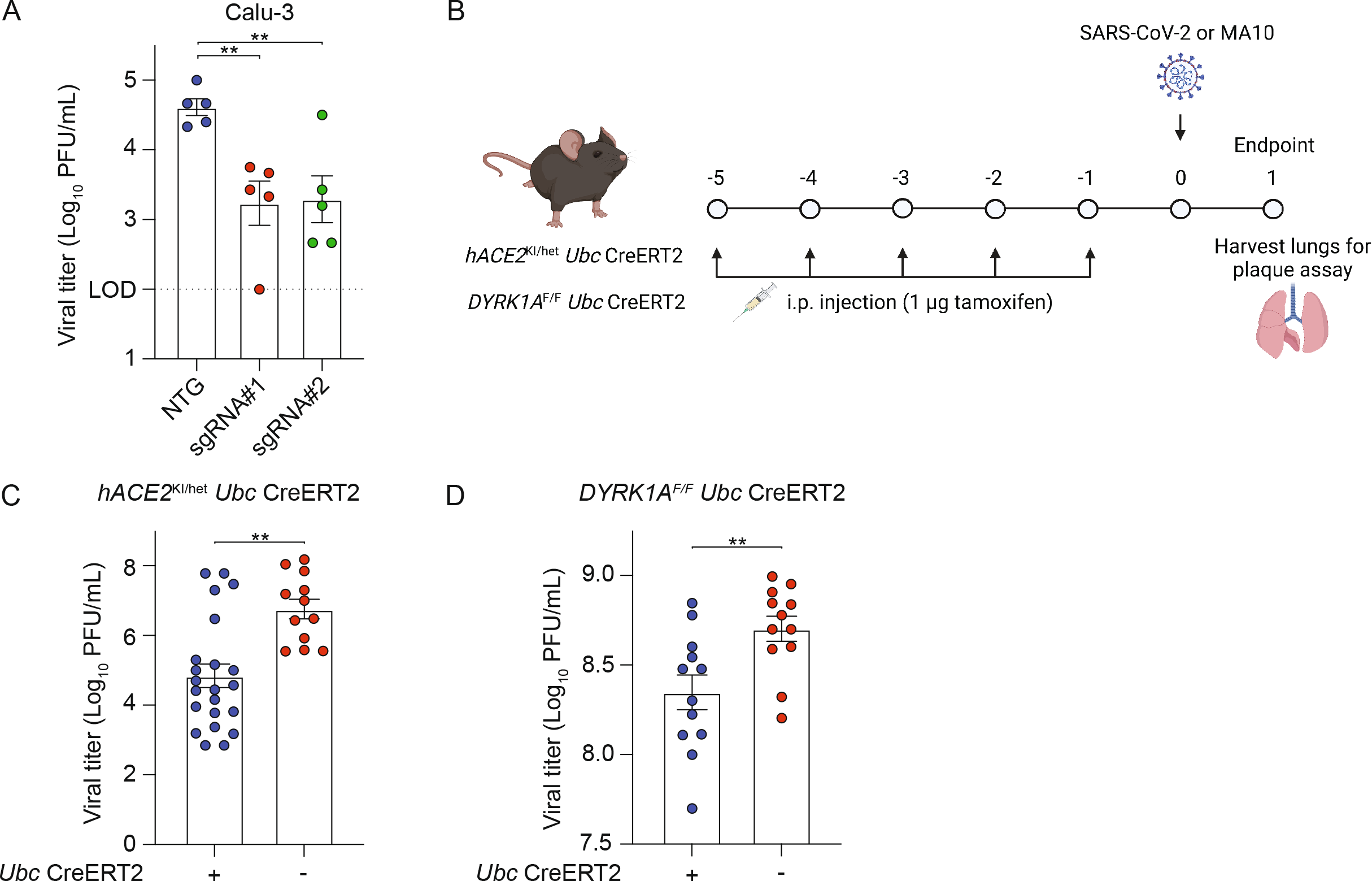
The proviral role of DYRK1A is conserved in human lung epithelial cells and in a murine model. **(A)** Polyclonal knockouts of DYRK1A were generated in Calu-3 human lung epithelial cells with two independent guides (sgRNA#1 and sgRNA#2). Viral titers were assessed after 24 hpi by TCID_50_ and compared against a non-targeting guide (NTG). **(B)** Conditional deletion and infection schematic for *hACE2^KI/het^ Ubc* CreERT2 and *DYRK1A^F/F^ Ubc* CreERT2. **(C)***ACE2* was conditionally deleted from a *hACE2* knock-in (KI) mouse under control of a tamoxifen inducible Cre recombinase (*hACE2^KI/het^ Ubc* CreERT2). Cre+ and Cre- mice were treated with tamoxifen for five days and were then infected intranasally with 10^5^ PFU SARS-CoV-2 (WA1/2020). **(D)***DYRK1A* was conditionally deleted from a *DYRK1AF/F* mouse (*DYRK1A^F/F^* Ubc CreERT2). Cre+ and Cre- mice were treated with tamoxifen for five days and were then infected intranasally with 10^5^ PFU MA10. In **(C, D)**, lung homogenates containing virus were harvested at 1-day post-infection. Lung viral titers were assessed by plaque assay and are reported as plaque forming units (PFU)/mL. Data shown are means ± SEM from **(A)** five, **(C)** three, or **(D)** two independent experiments. Each experiment in **(C, D)** included at least three mice per group. Data were analyzed by unpaired Student’s t-test; ** p< 0.01.

## DISCUSSION

During the COVID-19 pandemic, a number of genome-wide CRISPR/Cas9 screens have been performed to unveil host factors that regulate SARS-CoV-2 infection^22,23,27–33,91,92^. We previously identified *DYRK1A* as a pro-viral gene for both SARS-CoVs and MERS-CoV in Vero-E6 cells and Calu-3 cells^22,23^. Additional recent independent screens in Vero-E6 and Calu-3 cells have confirmed our initial finding of *DYRK1A* as a host dependency factor for SARS-CoV-2^33,34^. Numerous other SARS-CoV-2 genome-wide CRISPR knockout screens failed to identify DYRK1A – a disparity likely attributable to their reliance on cells that ectopically overexpress ACE2. In such cells (A549-ACE2 and Huh7.5-ACE2), ACE2 regulation is uncoupled from the transcriptional regulators that promote its endogenous expression, rendering them nonessential for coronavirus pathogenesis^27–30^. As a result, numerous transcription factors and epigenetic regulators are obscured in screening results. Here, we demonstrate that DYRK1A supports transcription of *ACE2* and *DPP4* by altering chromatin accessibility, dictating susceptibility to highly pathogenic human coronaviruses across various mammalian species.

Using single-cell knockout clones and addbacks, we reveal that nuclear DYRK1A promotes coronavirus entry by positively regulating *ACE2* and *DPP4* transcription via a kinase-independent mechanism. Wild-type and kinase-null (DYRK1A-Y321F) complementation *in vitro* offered significant but partial rescue of live SARS-CoV-2 infection (~70% of wild-type). Addbacks of another catalytically inactive mutant, DYRK1A-K188R, also exhibited partial but detectable rescue of SARS-CoV-2 infectivity. Of note, DYRK1A-K188R was not as effective at restoring icSARS-CoV-2-mNG infection relative to DYRK1A-Y321F. While this may result from minor residual kinase activity in DYRK1A-Y321F due to noncanonical autophosphorylation at Thr^111^, we anticipate this is likely not the case due to equal rescue of viral entry by DYRK1A-Y321F and DYRK1A-K188R in spike-dependent pseudovirus entry experiments^61,77,80^. Instead, we suggest this difference is explained by lower expression of DYRK1A-K188R due to previously described instability of the DYRK1A-K188R mutant^77^. Notably, DYRK1A-NES also partially restored viral entry, likely due to incomplete displacement of DYRK1A to the cytosol.

Highly effective vaccines and therapeutics like recombinant spike monoclonal antibodies, Paxlovid, remdesivir, and molnupiravir exist to mitigate COVID-19, but therapeutic targets that are conserved and cross-protective across highly pathogenic coronaviruses and variants are lacking^93–99^. A strengthened understanding of conserved coronavirus biology could reveal novel therapeutic targets and inform mechanistic ways to target them^100,101^. Although small molecule inhibitors of DYRK1A exist, these drugs are constrained by their limited selectivity despite ongoing improvements in their design^102,103^. Moreover, these inhibitors structurally target the catalytic function of DYRK1A, which we have shown to be dispensable for coronavirus pathogenesis. Small molecules such as PROTACs that could degrade DYRK1A may offer a novel therapeutic approach to combat coronavirus infection^104,105^. To date, no DYRK1A-specific PROTACs have been developed, but selective degraders of DYRK1A such as CaNDY have been developed^106^. It was also recently shown that SNX-544, an inhibitor of Hsp90, can greatly reduce SARS-CoV-2 infection in Vero-E6 and Calu-3 cells^107^. Inhibition of Hsp90 leads to destabilization and degradation of DYRK1A, suggesting that indirect mechanisms of DYRK1A depletion may exist that offer therapeutic benefit for SARS-CoV-2^106^. By conditionally deleting DYRK1A, we demonstrate in a small animal model that transient loss of DYRK1A is not only tolerable in adult small animals but also that it can reduce infection by SARS-CoV-2.

Our current study further shows that DYRK1A not only promotes expression of *ACE2* and *DPP4*, but also *CTSL*. Moreover, DYRK1A presence increases accessibility at previously identified putative *ACE2* regulatory elements (TSS, proximal enhancer, and a distal enhancer) and at sites that may regulate *DPP4* and *CTSL* (Wei and Patil *et al*. 2022). As receptor and protease expression are critical for coronavirus entry, these findings suggest that DYRK1A operates as a critical modulator of chromatin accessibility and transcription to drive a pro-viral gene program that promotes viral entry. However, deletion of DYRK1A in Calu-3 cells – which lowly express CTSL and instead employ TMPRSS2 for spike proteolytic processing – still conferred resistance to SARS-CoV-2 infection. Nonetheless, we cannot preclude the possibility that DYRK1A may also regulate TMPRSS2 and other aspects of viral entry besides receptor expression,

DYRK1A was previously reported to function as a positive regulator of transcriptional activity^58,64,65,67,68^. In the nucleus, DYRK1A can be recruited to enhancers and promoter regions, facilitating formation of the preinitiation complex or chromatin accessibility^64,65,67,108,109^. DYRK1A does not contain a DNA-binding domain and can bind and/or phosphorylate other proteins at these sites, supporting the function of other proteins such as RNA polymerase II (RNAPII) and histone acetyltransferases (P300/CBP)^65,108^. DYRK1A can also potentiate transcription independently of its kinase activity, such as by recruiting RNAPII to form the pre-initiation complex or by binding known transcription factors as a scaffold, like androgen receptor-interacting protein 4 (ARIP4, also known as RAD54L2) and forkhead box O1 (FOXO1)^64,65,67^. Among the top pro-viral genes identified in our original CRISPR screen, many genes encode known DYRK1A interactors involved in transcriptional regulation, including those coding for the SWI/SNF complex (*SMARCA4, SMARCB1, ARID1A, SMARCE1, SMARCC1, DPF2)*, a histone methylase/demethylase complex (*KMT2D, KDM6A*), and transcriptional coregulators (*ARIP4/RAD54L2*)^22,39,58,110^.

Because SMARCA4, KDM6A, and HMGB1 are known *ACE2* regulators, we asked whether DYRK1A may cooperate with one or more of these proteins, serving as a scaffold to promote *ACE2* or *DPP4* gene transcription^22,111^(Wei and Patil *et al*. 2022). Both HMGB1 and SMARCA4 alter chromatin accessibility at the *ACE2* locus, but how KDM6A modulates *ACE2* expression has yet to be defined ^22^(Wei and Patil *et al*. 2022). DYRK1A-HMGB1 interactions have not been described, but DYRK1A can interact with SMARCA4 (the catalytic subunit of SWI/SNF) and SMARCB1 (a core subunit of SWI/SNF that stabilizes it and targets it to enhancers), resulting in phosphorylation of SMARCB1^58,110,112^. Whether phosphorylation of SMARCB1 is critical for SWI/SNF activity is unclear, but SMARCA4 can rearrange nucleosomes at the *ACE2* promoter and three putative enhancers, driving accessibility and transcription of *ACE2*^113^ (Wei and Patil *et al*. 2022). Overall, the DEGs and chromatin regions regulated by DYRK1A poorly correlate with those altered by HMGB1 or SMARCA4, although some pathways are shared between DYRK1A and SMARCA4 (**Extended Data Fig. 3**)^22^(Wei and Patil *et al*. 2022). Furthermore, HMGB1 and SMARCA4 do not alter *DPP4* expression or promote MERS-CoV infection^22^. Together these findings suggest that DYRK1A likely functions independently of HMGB1 and SWI/SNF to promote CoV infection. However, we cannot exclude the possibility that SMARCA4 and DYRK1A may coordinate to co-regulate some sites (i.e., *ACE2*) but not others (i.e., *DPP4*). Unlike SMARCA4 and HMGB1, the histone methyltransferase KDM6A regulates both SARS-CoV-2 and MERS-CoV infection^22,111^(Wei and Patil *et al*. 2022). KDM6A complexes with KMT2D, a histone demethylase, and the H3K27 acetyltransferase P300 and can activate enhancers^114^. DYRK1A can directly interact with KMT2D and P300, hyper-phosphorylating P300 to drive H3K27 deposition and enhancer accessibility^39,108^. Because kinase activity is dispensable for DYRK1A-driven *ACE2* and *DPP4* expression, we suspect that DYRK1A also operates independently of KDM6A and KMT2D via a novel kinase-independent mechanism but cannot preclude that DYRK1A may act as a scaffold for these proteins. Clarifying the proteins that coordinate with DYRK1A, including transcription factor(s) that target DYRK1A to these sites, represent an important future direction of this work.

ACE2 and DPP4 perform physiologic functions independent of their roles as coronavirus receptors. *ACE2* encodes a dipeptidyl carboxypeptidase that cleaves angiotensin I, maintaining homeostasis in the renin-angiotensin system^115^. Located on chromosome *Xp22*, ACE2 is an X-linked, dosage-sensitive gene known to be transcriptionally regulated by EP300, SP-1, CEBP, GATA3, and HNF1/4^115,116^. Recent efforts have aimed to identify novel ACE2 modifiers, uncovering *PIAS1, SMAD4, BAMBI*, *KDM6A*, and *GATA6*^111,117^. *DPP4*, which is situated on chromosome *2q24, e*ncodes a serine exopeptidase that regulates glucose homeostasis^118,119^. The *DPP4* promoter contains binding sites for NF-kB, SP-1, EGFR, NF-1, STAT1, and HNF1^120–122^. Until now, only hepatocyte nuclear factor (HNF) family proteins have been identified as shared transcriptional regulators for these genes. Here, we demonstrate for the first time that DYRK1A is a novel regulator for both genes. However, the impact of DYRK1A on the renin-angiotensin system and glucose homeostasis have yet to be thoroughly explored and warrant additional investigation. DYRK1A is also highly dosage-sensitive, and whether fine-tuning of DYRK1A expression could alter these critical biological pathways is unknown.

In Down syndrome (trisomy 21), DYRK1A is overexpressed 1.5-fold due to its position on chromosome 21, and dysregulation of DYRK1A dosage directly contributes to neurological defects and disease pathogenesis^40–47^. Strikingly, Down syndrome significantly increases (5-10x) the risk of COVID-19 infection, hospitalization, and death relative to the general population^49–54^. While individuals with Down syndrome possess many comorbidities that could partially explain their predisposition to COVID-19 severity, together they incompletely explain such an elevated risk. A recent network analysis revealed that *TMPRSS2* – which is also located on chromosome 21 and is subsequently overexpressed in Down syndrome – may contribute to COVID-19 severity by promoting increased viral entry^50,52^. Unlike TMPRSS2, DYRK1A does not directly interact with SARS-CoV-2, but we now show that DYRK1A positively regulates ACE2 expression and viral entry. Although *ACE2* is not globally overexpressed in Down syndrome, few tissues have detectable ACE2 expression at baseline and transcriptomic datasets for airway epithelial cells derived from trisomy 21 are lacking^50^. Whether DYRK1A overexpression in trisomy 21 is sufficient to elevate ACE2 expression and contribute to COVID-19 severity remains to be seen and warrants further investigation.

The role of DYRK1A in Down syndrome has been extensively studied, but its function in coronavirus biology had not previously been elucidated. Our findings now support that DYRK1A can promote DNA accessibility independently of its kinase activity, facilitating ACE2 and DPP4 expression for viral entry. We reveal that the pro-viral function of DYRK1A dictates susceptibility to diverse highly pathogenic human coronaviruses and is conserved across multiple mammalian species. Thus, DYRK1A represents a novel therapeutic target for multiple coronavirus lineages. While chemically targeting DYRK1A independently of its kinase activity remains challenging, our work highlights the value of considering genetic regulators or interactors in informed therapeutic target design.

## MATERIALS AND METHODS

### Cell culture

HEK293T (ATCC) and Vero-E6 (ATCC) cells were cultured in Dulbecco’s Modified Eagle Medium (DMEM, Gibco) with 5% heat-inactivated fetal bovine serum (FBS, VWR) and 1% Penicillin/Streptomycin (Gibco) unless otherwise noted. Vero-E6-ACE2-TMPRSS2 (gift from Barney Graham, NIH) were cultured with 5% DMEM, 5% FBS, 1% Penicillin/Streptomycin, 5 μg/ml puromycin, and 5 μg/ml blasticidin. Calu-3 (ATCC) cells were cultured in RPMI 1640 (Gibco) with 1% Glutamax 100X (Gibco), 10% FBS, 1% Penicillin/Streptomycin and 16 ng/ml hepatocyte growth factor (HGF, Stem Cell Technologies). When selecting cells transduced by lentivirus, Vero-E6 cells and Calu-3 were treated with 5 μg/ml or 1 μg/ml puromycin (Gibco), respectively. All cells were grown at 37°C/5% CO_2_.

### Expression constructs, lentiviral packaging, and lentiviral transduction

All constructs were PCR amplified from a codon optimized gene block encoding the coding sequence of human DYRK1A (GenScript) using Q5 High-Fidelity DNA Polymerase with GC enhancer buffer (New England Biolabs). DYRK1A was cloned by Gibson assembly into pCW57.1-puro (gift of Katerina Politi, Yale School of Medicine). pCW57.1-DYRK1A K188R, Y321F, and NES constructs were generated by Q5 site-directed mutagenesis according to manufacturer instructions (New England Biolabs). All constructs were sequence validated by Sanger sequencing or Plasmidsaurus Oxford Nanopore sequencing. To generate lentiviral particles, a three-component lentiviral system using 2.4 ug pVSV-G, 4.8 ug pSPAX2, and 8 ug lentiviral plasmid via calcium phosphate transfection of 50-70% confluent HEK293T in a 10 cm dish. Supernatant containing lentivirus was harvested for three consecutive days and pooled. Cellular debris was clarified by centrifugation at 500 xg/5 min. For complementation, lentivirus was concentrated approximately 50-fold by using 4x PEGit lentiviral concentrator (MD Anderson). For lentiviral transduction, Vero-E6 or Calu-3 cells were transduced at 50% confluency and selected with puromycin 48 hours later. hACE2 and hDPP4 overexpressing lines were generated by stable lentiviral delivery of pLV-EF1a-ACE2-puro (gift of A. Iwasaki) and pLEX307-DPP4-puro (Addgene #158451) into *DYRK1A* knockout clones.

### Viral stocks

Viral stocks were generated in Vero-E6 or Vero-E6-ACE2-TMPRSS2 cells seeded at ~80% confluency inoculated with HKU5-SARS-CoV-1-S (NR-48814), SARS-CoV-2 isolate USA-WA1/2020 (NR-52281), Germany isolate B (NR-52370), SARS-CoV-2 isolate B.1.1.529 (NR-56481), or MERS-CoV (icMERS-CoV EMC/2012) (NR-48813) from BEI resources at a MOI of approximately 0.01 for three days to generate a P1 stock. Vero-E6 or Vero-E6-ACE2-TMPRSS2 cells were inoculated with the P1 stock and incubated for three days or until 50-70% cytopathic effect was observed to generate a P2 stock. To generate icSARS-CoV-2-mNG stocks, lyophilized icSARS-CoV-2-mNG (World Reference enter for Emerging Viruses and Arboviruses, Galveston, TX) was resuspended in 500 μl deionized water and diluted 100-fold in medium. Diluted virus was added to 10^7^ Vero-E6 cells grown in T175 (Corning) for three days. Vero-E6-ACE2-TMPRSS2 cells at >50% confluency were inoculated with 10^6^ PFU of ic2019-nCoV MA10 (gift from Ralph Baric, University of North Carolina Chapel Hill) in 2 mL 1X PBS and incubated for 1 hour at 37°C with periodic rocking to facilitate viral adherence. One hour later, fresh medium was added to the cells and incubated overnight. Near complete cytopathic effect was observed at 1 dpi, and viral supernatant was collected. All viral stock harvests were clarified by centrifugation (500 xg/5 min) and filtered through a 0.45 μm filter (Millipore Sigma) aliquoted, and stored at −80°C. All viral stocks were tittered by at least two independent plaque assays or TCID_50_ assays. Final viral stocks generated at Yale were sequenced by the laboratory of Nathan Grubaugh (Yale University School of Public Health) to confirm no substitutions were generated during viral stock propagation. All work with infectious virus was performed in a Biosafety Level 3 facility in accordance to regulations and approval from the Yale University Biosafety Committee and Yale University Environmental Health and Safety.

### SARS-CoV-2 plaque assays

Vero-E6 cells were seeded at 4×10^5^ cells/well in 12-well plates (Corning) and were incubated overnight. The next day, media were removed and replaced with 100 μl of 10-fold serial dilutions of virus. Plates were incubated at 37C for 1 hour, with gently rocking every 15 minutes to promote viral adherence. Wells were then covered with 1 mL overlay media (DMEM, 2% FBS, 0.6% Avicel RC-581) and incubated for 48 hours at 37C. At 2dpi, plates were fixed with 10% formaldehyde (Ricca Chemical) for 30 min then stained with crystal violet solution (0.5% crystal violet (Sigma-Aldrich) in 20% ethanol) for 30 min. Crystal violet was aspirated and wells were rinsed with deionized water to visualize plaques.

### SARS-CoV-2 infections of polyclonal Calu-3 cells by TCID_50_

Polyclonal knockouts of DYRK1A were generated in Calu-3 cells using two independent guides (sgRNA#1 and sgRNA#2). Calu-3 cells were seeded into 24-well plates at a density of 2 × 10^5^ cells/well. Two days later, cells were infected with SARS-CoV-2 post-seeding at an MOI ~ 0.05. Viral inocula were incubated with cells for 30 min at 37 °C. Unbound virus was aspirated and cells were washed once with 1X PBS. Cells were incubated for 24 hours and then subjected to mechanical lysis by freeze thaw. Infectious viral particles were tittered by serial dilution and incubated on 96-well plates coated with Vero-E6 cells. Each dilution was applied to eight wells. Cytopathic effect was determined visually at 3 dpi, and TCID_50_/mL was calculated using the dilution factor required to produce CPE in half of the wells. Viral titers were assessed and compared against the non-targeting guide (NTG).

### SARS-CoV-2 fluorescent reporter virus assay

Cells were plated at 2500 cells/well in a 384-well plate (Greiner) and adhered at 37C for approximately 5 hours. icSARS-CoV-2-mNG was added at an MOI of 1.0. Infection frequency was measured by mNeonGreen expression at 2dpi by high content imaging (Cytation 5, BioTek) configured with bright field and GFP cubes. Total cell numbers were determined from bright field images using Gen5 software. Object analysis was used to quantify the number of mNeonGreen positive cells. Percent infection was calculated as the ratio between mNeonGreen+ cells and total cells.

### Generation of DYRK1A clonal Vero-E6 knockout and complemented cells

Vero-E6 DYRK1A KO cells were generated by lipofection of Cas9-Ribonucleoproteins (RNPs). CRISPR guide RNAs (gRNA) were synthesized by IDT (sequence: TCAGCAACCTCTAACCAACC). gRNAs were complexed in a 1:1 molar ratio with tracrRNA in nuclease-free duplex buffer by heating at 95°C for 5 min and then cooled to room temperature. Duplexes were combined with Alt-R Cas9 enzyme at room temperature for 5 min to form RNPs in Opti-MEM with 200 μl total volume. Complexes were mixed with RNAiMAX transfection reagent (Invitrogen) in Opti-MEM at room temperature for 20 min before transfection. Transfection was performed with 3.2 × 10^5^ Vero-E6 cells in suspension in a 12-well plate (Corning). Cells were incubated for 48 hours then stained with 1:500 Zombie Aqua (BioLegend) in 1X PBS (Gibco) prior to flow cytometry-based sorting on live cells. Single cells were sorted into 96-well plates (Corning). Clones were screened by resistance to rcVSV-SARS2-S virus-induced cell death, Western blot, and Sanger sequencing. DYRK1A KO clones were complemented by lentiviral transduction of pCW57.1-puro vector containing full-length DYRK1A, kinase-dead DYRK1A (K188R or Y321F), or nuclear localization defective DYRK1A (NES) with an N-terminal 3X FLAG tag. Two days post-transduction, puromycin was added and cells were selected for three days to select for stably expressing addbacks. Stable DYRK1A expression was induced by the addition of 20 or 200 ug/ml doxycycline hyclate (Sigma-Aldrich) dissolved in DMSO (Sigma-Aldrich) for 24 or 72 hours. Expression of DYRK1A in complemented cells was confirmed by Western blot for 3X FLAG and DYRK1A.

### Generation of DYRK1A polyclonal Calu-3 knockout cells

Oligonucleotides (Yale, Keck Oligo) were generated with BsmBI-compatible overhangs (guide 1 pair: CACCGTCAGCAACCTCTAACTAACC, AAACGGTTAGTTAGAGGTTGCTGAC; guide 2 pair: CACCGTGAGAAACACCAATTTCCGA, AAACTCGGAAATTGGTGTTTCTCAC). Oligos were annealed and phosphorylated using equimolar ratios of oligo pairs with 1X T4 Ligation Buffer (New England Biolabs) and T4 PNK (New England Biolabs) at 37°C for 30 min, 95°C for 5 min, then −5°C/min to 25°C. LentiCRISPRv2 (Addgene #52961) was digested by BsmBI-v2 (New England Biolabs) for 2 hours at 55°C in 1X NEBuffer 3.1. Double-stranded oligonucleotides were ligated into the digested lentiCRISPRv2 vector using T4 DNA ligase (New England Biolabs) according to manufacturer protocol. Ligated vectors were transformed into Stbl3 cells and sequence verified for correct guide insertion. Lentiviral plasmids were co-transfected with packaging plasmids pSPAX2 and pVSV-G in HEK293T cells and Calu-3 cells were transduced with lentiviral particles. Transduced cells were selected with puromycin for two weeks prior to infection.

### Pseudovirus production

VSV-based pseudotype viruses were generated as previously described in 10 cm dishes^123^. Briefly, HEK293T cells were transfected with pCAGGS or pCDNA3.1 vectors expressing the CoV spike (S) glycoprotein by calcium phosphate transfection and then inoculated with a replication-deficient VSV encoding Renilla luciferase in place of the G glycoprotein. After one hour at 37°C, unbound inoculum was removed and cells were washed with 4X PBS. Fresh media were added with anti-VSV-G clone I1 (8G5F11) (Absolute Antibody) to neutralize residual VSV-G virus^124^. After 24 hours, supernatant was harvested as viral stock and centrifuged at 3000 rpm for 10 min to clarify cellular debris. VSVpp-SARS2-S stocks were concentrated using Amicon Ultra 100 kD filter columns (Millipore Sigma) at 3000 rpm. VSVpp-SARS1-S and VSVpp-MERS-S stocks were not concentrated. All stocks were aliquoted and stored at −80°C. Plasmids encoding codon-optimized sequences of SARS-CoV-S and MERS-SΔCT were previously described^11,125^. Vector pCAGSS containing the SARS-CoV-2, Wuhan-Hu-1 S Glycoprotein Gene (NR-52310) was produced under HHSN272201400008C and obtained through BEI Resources, NIAID, NIH.

### Pseudovirus entry assay

4 × 10^5^ Vero-E6 cells were seeded in 100 μl volume of each well of a black-walled clear bottom 96-well plate (Corning) and incubated at 37°C for approximately 5 hours to allow cells to adhere. VSV pseudovirus at 1:10 final concentration volume/volume for unconcentrated virus (VSVpp-SARS1-S or VSVpp-MERS-S) or 1:20 for concentrated virus (VSVpp-SARS2-S). Virus was incubated with cells for 24 hours and cells were subsequently lysed with Renilla Luciferase Assay System (Promega) according to manufacturer instructions. Luciferase activity was measured using a microplate reader (BioTek Synergy). Pseudovirus entry was normalized to VSV-G within each condition, and percent entry was calculated relative to wild-type Vero-E6 cells after normalization.

### RT-qPCR

Total RNA was isolated from cells and lung homogenates using Direct-zol RNA MiniPrep Plus kit (Zymo Research) and 500 ng RNA was used for cDNA synthesis. Quantitative PCR was carried out using 2 μl cDNA (diluted 1:10) with specific primers and probes (IDT) for *b-actin* (Forward: 5’-GGATCAGCAAGCAGGAGTATG-3’; Reverse: 5’-AGAAAGGGTGTAACGCAACTAA-3’; Probe: /56-FAM/TCGTCCACC/ZEN/GCAAATGCTTCTAGG/3IABkFQ/), *ACE2* (Forward: 5’-AGAGGATCAGGAGTTGACATAGA-3’; Reverse: 5’-ACTTGGGTTGGGCACTATTC-3’; Probe: /56-FAM/ACCGTGTGG/ZEN/AGGCTTTCTTACTTCC/3IABkFQ/), and *DPP4* (Forward: 5’-GACATGGGCAACACAAGAAAG-3’; Reverse: 5’-GCCACTAAGCAGTTCCATCT-3’; Probe: /56-FAM/TTTGCAGTG/ZEN/GCTCAGGAGGATTCA/3IABkFQ/) genes. Reactions were prepared according to manufacturer recommendations for AmpliTaq Gold DNA polymerase (Applied Biosystems).

### Western blotting

1×10^6^ cells were collected and lysed in Alfa Aesar Nonidet 40 (NP-40; 20 mM Tris-Hcl [pH 7.4], 150 mM NaCl, 1 mM EDTA, 1% Nonidet P-40, 10 mg/ml aprotinin, 10 mg/ml peupeptin, and 1 mM PMSF). Cell lysates were fractionated on SDS-PAGE pre-cast gels (BioRad) and transferred to a PVDF membrane by TurboTransfer (BioRad). Immunoblotting assays were performed with the following primary antibodies (1:1000): anti-ACE2 (ProSci, cat#3217), anti-DYRK1A (Abcam, cat#ab259869), anti-FLAG (Sigma-Aldrich, cat#F3165), anti-DPP4 (R&D, cat#AF1180), anti-GAPDH (BioLegend, cat#649202), anti-Lamin B1 (BioLegend, cat# 869801). Proteins were visualized with goat anti-mouse or goat anti-rabbit IgG secondary antibodies (1:5000) diluted in 2% Omniblot milk (AmericanBio) in 1X TBST using a chemiluminescence detection system (BioRad ChemiDoc MP).

### SARS-CoV-2 *in vitro* DYRK1A inhibition assays

Harmine and INDY were purchased from Cayman Chemical, and DYR219 was synthesized in-house. Compounds were resuspended at a stock concentration of 40-50 mM in DMSO. Drugs were diluted 2-fold in DMSO and spotted into 384-well black skirted plates (Corning) in 20 nL at 1000X drug stock using the Labcyte ECHO dispenser at the Yale Center for Molecular Drug Discovery. 1.25×10^3^ Vero-E6 cells were plated in total volume of 20 μl. Two days later, cells were infected with MOI ~ 1 SARS-CoV-2 isolate USA-WA1/2020 in 5 μl media. Cells were incubated for three days before assessing viability and virus-induced cell death by CellTiter Glo according to manufacturer protocol (Promega). Luminescence was quantified using a plate reader (Cytation 5, BioTek). For each cell line, viability was determined in SARS-CoV-2 infected cells relative to uninfected cells.

### RNA-seq

WT Vero-E6 cells, DYRK1A KO#1 + vector, and DYRK1A KO#1 + complements were seeded at 3 × 10^5^ cells/well in a 6-well plate and were treated with doxycycline for 72 hours to enable rescue of *ACE2* and *DPP4*. Samples were performed in biological duplicate and harvested by scraping. Total cellular RNA was extracted using the Direct-zol RNA MiniPrep Plus (Zymo Research) and libraries were prepared with rRNA depletion by the Yale Center for Genome Analysis. RNA-seq libraries were sequenced on Illumina NovaSeq 6000 with a goal of at least 25 × 10^6^ reads per sample. Reads were aligned to reference genome chlSab2, NCBI annotation release 100 using STAR aligner v2.7.3a with parameters –winAnchorMultimapNmax 200 – outFilterMultimapNmax 100 –quantMode GeneCounts^126^. Differential gene expression was obtained using the R package DESeq2 v1.32^127^. Heatmaps were generated using R package^128^.

### ATAC-seq

WT Vero-E6 cells, DYRK1A KO#1 + vector, and DYRK1A KO#1 + complement were seeded at 3 × 10^5^ cells/well in a 6-well plate and were treated with doxycycline for 72 hours to enable rescue of *ACE2* and *DPP4*. Samples were performed in biological duplicate and were harvested by scraping. Samples were submitted to Yale Center for Genome Analysis for library generation and were sequenced on an Illumina NovaSeq S4 instrument as 101 nt long paired-end reads with goal of at least 45 × 10^5^ reads per replicate. Reads were trimmed of Nextera adaptor sequences using Trimmomatic v0.39^129^ and aligned to chlSab2 using Bowtie2 v2.2.9^130^ with parameter –X2000. Duplicates were marked using Picard Tools v2.9.0 (Broad Institute. version 2.9.0. “Picard Tools.” Broad Institute, GitHub repository. http://broadinstitute.github.io/picard/). Duplicated, unpaired, and mitochondrial reads were removed using SAMTools v1.9^131^. Reads were shifted +4 bp and −5 bp for forward and reverse strands, respectively. Peaks were called using MACS2 v2.2.6^132^ with parameters –nomodel – keep-dup all -s 1 –shift 75 –extsize 150. Reads that fell inside peaks were counted using featureCounts v1.6.2^133^ and differential accessibility analysis was performed using DESeq2 v1.32^127^. Bigwig files were generated using deeptools v3.1.3 with parameter –normalizeUsing RPKM^134^. Data were visualized with Integrated Genome Viewer.

### Generation of DYRK1A conditional knockout mice

*DYRK1A^F/F^* mice were obtained from the Jackson Laboratory (Strain #027801) and crossed to *Ubc*-Cre-ERT2 mice (Strain #007001). Mice were genotyped using primer and probe sequences provided by Transnetyx, Inc. (*DYRK1A* Flox: Forward: 5’-TGTATGCTATACGAAGTTATTAGGTCCCT-3’, Reverse: 5’-CTTTTGTTAGTGTATGGCATAACTTGCA-3’, Reporter (FAM): 5’-CAGTGGGAGGATCCCCT-3’; *Ubc*-Cre-ERT2: Forward: 5’-AGGGCGCGCCGAATT-3’, Reverse: 5’-GGTAATGCAGGCAAATTTTGGTGTA-3’, Reporter (FAM): 5’-CCACCATGTCCAATTTA-3’.

### Mouse infections

Sex-matched, age-matched, litter-mate controls were used for all experiments. *DYRK1A^F/F^*, *Ubc* CreERT2, and *hACE2* knock-in (KI) mice were obtained from the Jackson Laboratory and crossed in-house. To activate the Cre recombinase and conditionally delete DYRK1A, *DYRK1A^F/F^ Ubc* CreERT2 or *hACE2* KI/het *Ubc* CreERT2 *+/−* and *−/−* mice were treated with 1 μg tamoxifen in corn oil (Sigma) per day for five consecutive days. On day six, mice were then infected with 5 × 10^5^ PFU intranasally with MA10 or 1 × 10^5^ PFU SARS-CoV-2 WA/01/2020 or icSARS-CoV-2 WA/01/2020. PFU for infection were calculated by plaque assay on WT Vero-E6 cells. Mice were sacrificed at 1 dpi and lungs were collected in 1 mL DMEM/2% FBS/1X Antibiotic-Antimycotic (Gibco). Samples were homogenized and debris was clarified at maximum speed for 10 minutes at 4°C. Lung homogenate was aliquoted and stored at −80°C for plaque assays. Remaining lung homogenate was mixed 1:3 with TRIzol LS for one hour at room temperature to inactivate virus. RNA extraction was performed on inactivated virus with Direct-zol RNA MiniPrep Plus according to manufacturer instructions. *DYRK1A* gene disruption and *ACE2* reduction were quantified by RT-qPCR.

### Ethics statement

The care and use of all animals were approved in accordance with the Yale Animal Resource Center and Institution Animal Care and Use Committee (#2021-20198) in agreement with the standards set by the *Animal Welfare Act*.

### Statistical analysis

All statistical analysis was performed in Prism GraphPad version 9.2.0 (San Diego, CA). Error bars indicate standard error of the mean unless otherwise indicated. Normally distributed data was analyzed using unpaired Student’s t-tests while Mann-Whitney tests were performed for non-normally distributed data. For more than two comparisons, ordinary one-way ANOVA or Kruskal Wallis tests were performed according to normality. *P* values of <0.05 were considered statistically significant (*, *P* < 0.05; **, *P* < 0.01; ***, *P* < 0.001; ****, *P* < 0.0001).

## Data availability

GEO accession# pending. All mice are available for purchase at the Jackson Laboratories. Viral stocks and plasmids are available under Material Transfer Agreement.

## Acknowledgements

We would like to acknowledge the Yale Center for Molecular Discovery, Yale Center for Genome Analysis, and the Yale Flow Cytometry Facility for technical assistance and sample processing, and BEI Resources, the World Reference Center for Emerging Viruses and Arboviruses (WRECVA), Moitrayee Bhattacharyya, Akiko Iwasaki, Katerina Politi, Yoshihiko Miyata, Man Mohan, and Vance Lemmon for critical reagents and expertise.

## Author contributions

M.S.S., J.W., and C.B.W conceptualized the study and designed the experiments. M.S.S, W.L.C., J.W., M.M.A., A.P., K.S.C., C.K.C., R.B.F., S.B.B, P.H., P.C.D., R.H., and K.S. performed the research. M.S.S., W.L.C., J.W., A.P., K.S.C., C.K.C., P.C.D., and R.H. analyzed the data. M.S.S. wrote the paper, and all authors provided input. P.H., C.H., S.V., J.G.D., C.K., Q.Y., and C.B.W. provided resources and funding acquisition.

