## Extended Figures for "Kinase-independent activity of DYRK1A promotes viral entry of highly pathogenic human coronaviruses"

### Extended Data Figure 1

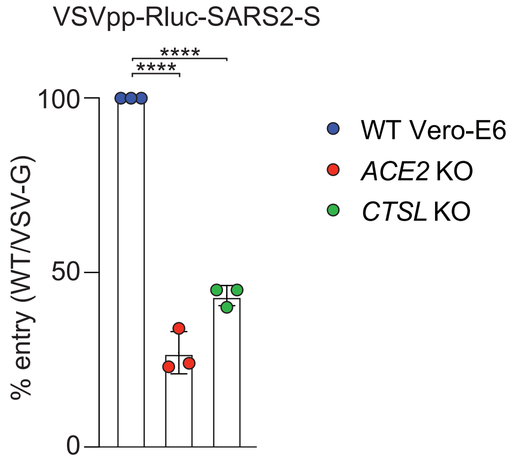


**Extended Data Figure 1. Loss of ACE2 and CTSL reduce SARS-CoV-2 pseudovirus entry.** Polyclonal deletion of *ACE2* or *CTSL* significantly reduce pseudotyped particles expressing the SARS-CoV-2 spike relative to WT Vero-E6 cells, like loss of *DYRK1A*. Cells were infected with VSV pseudovirus (VSVpp) encoding CoV spike proteins and a Renilla luciferase (Rluc) reporter at 24 hpi. % Entry for VSVpp-Rluc-SARS2-S was normalized to VSVpp-Rluc-VSV-G control and WT Vero-E6 cells. Data were analyzed by unpaired Student's t-test; **** p< 0.001. Shown are means ± SEM. Data are representative of three independent biological experiments performed with 3 technical replicates.

### Extended Data Figure 2

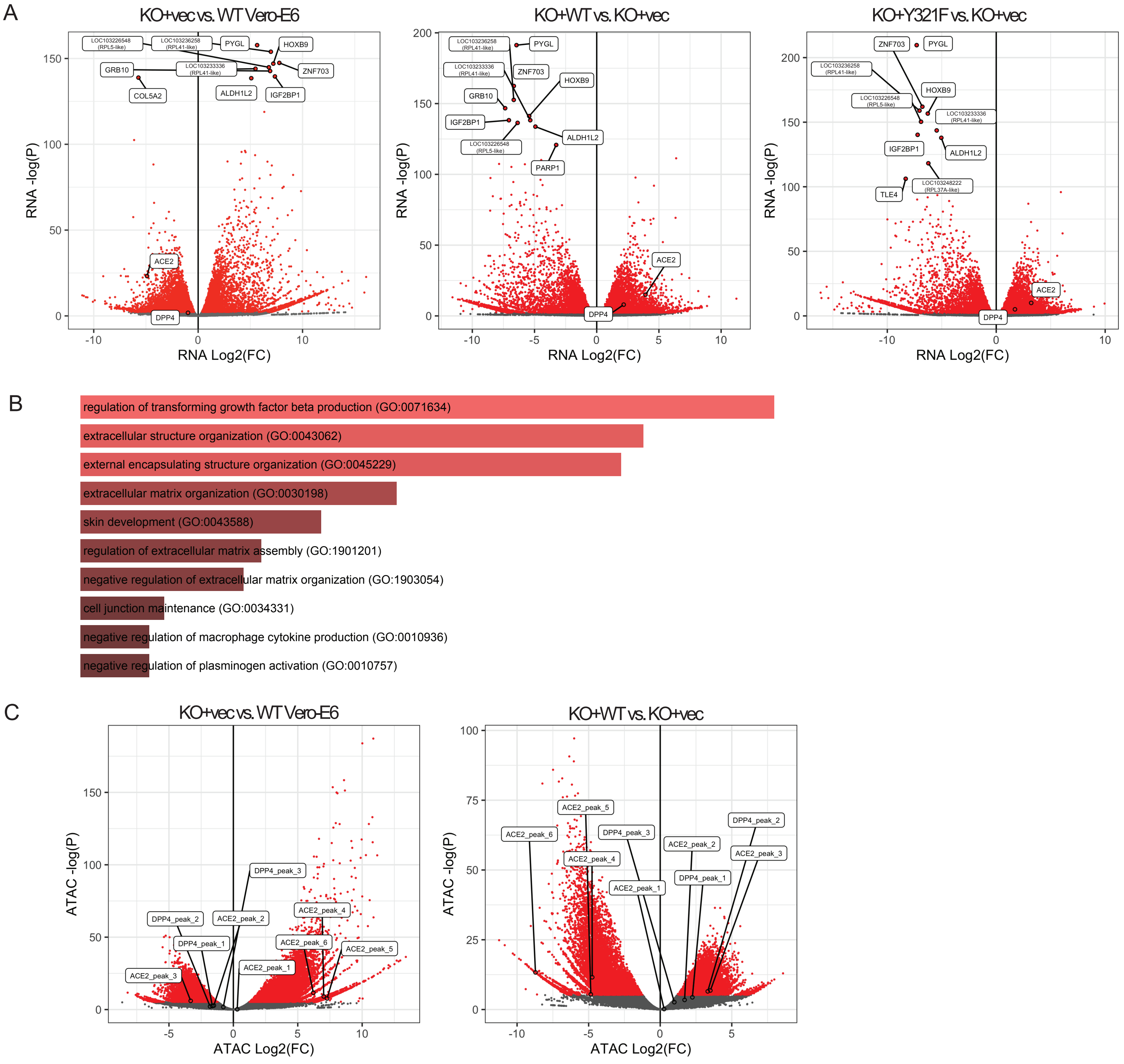


**Extended Data Figure 2. DYRK1A regulates global gene expression and chromatin accessibility. (A)** RNA-seq volcano plots depicting differentially expressed genes in cells where DYRK1A is absent or reintroduced. **(B)** Enrichr pathway analysis for biological function enriched in the presence of DYRK1A. Gene set criteria included p<0.05, L2FC<0 for KO+vec vs. WT Vero-E6 and L2FC>1.5 for both KO+WT and KO+Y321F vs. KO+vec. **(C)** ATAC-seq volcano plots depicting differentially expressed genes in cells where DYRK1A is absent or reintroduced. All experiments were performed in biological duplicate.

### Extended Data Figure 3

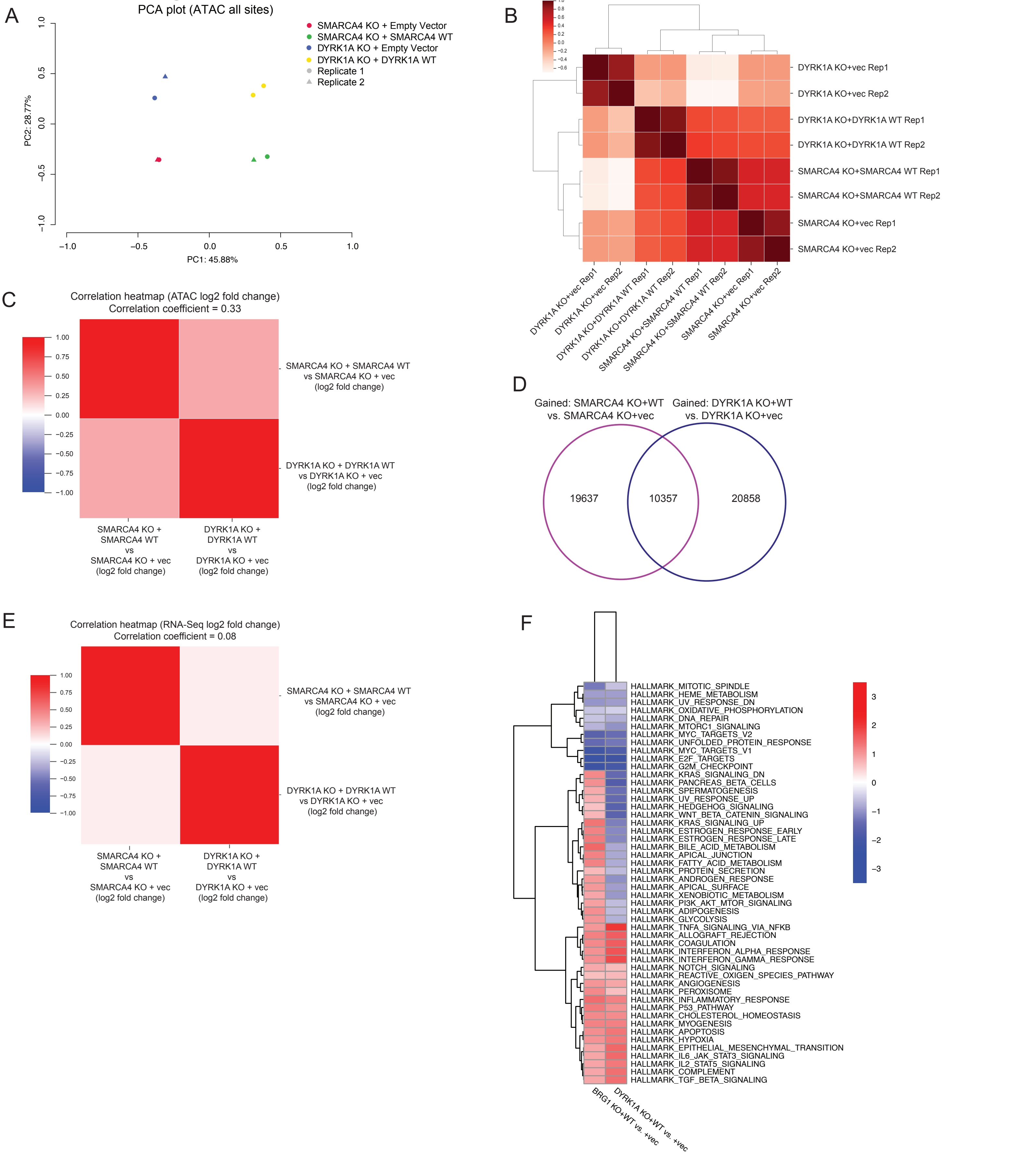


**Extended Data Figure 3. DYRK1A and SMARCA4 share limited similarities in regulating chromatin accessibility and gene expression. (A)** Principal component analysis of ATAC-Seq experiments performed in DYRK1A KO+vec, DYRK1A KO+WT, SMARCA4 KO+vec, and SMARCA4 KO+WT cells generated in a parental Vero-E6 background. Each experiment was performed in biological duplicate (replicate 1 and 2). DYRK1A and SMARCA4 loss share some molecular impacts, suggesting some pathways may be co-regulated (PC1) whereas others may be independently regulated (PC2). **(B)** Correlation heatmap comparing all sites from ATAC-Seq experiments in DYRK1A KO+vec, DYRK1A KO+WT, SMARCA4 KO+vec, and SMARCA4 KO+WT cells. **(C)** Correlation heatmap comparing chromatin accessibility by ATAC-Seq in DYRK1A or SMARCA4 complemented cells, identifying a correlation coefficient of 0.33 supporting ~33% of clusters may be corelated by DYRK1A and SMARCA4. **(D)** Venn diagram highlighting shared peaks gained by DYRK1A and SMARCA4 complementation. **(E)** Correlation heatmap comparing changes in RNA abundance in DYRK1A or SMARCA4 complemented cells, identifying a correlation coefficient of 0.08 supporting <10% of the top upregulated/downregulated genes are shared between DYRK1A and SMARCA4. **(F)** Gene set enrichment analysis from RNA-Seq experiments showing shared pathway regulation by DYRK1A and SMARCA4.
